# DeepARG: A deep learning approach for predicting antibiotic resistance genes from metagenomic data

**DOI:** 10.1101/149328

**Authors:** G. A. Arango-Argoty, E. Garner, A. Pruden, L. S. Heath, P. Vikesland, L. Zhang

## Abstract

Growing concerns regarding increasing rates of antibiotic resistance call for global monitoring efforts. Monitoring of environmental media (e.g., wastewater, agricultural waste, food, and water) is of particular interest as these media can serve as sources of potential novel antibiotic resistance genes (ARGs), as hot spots for ARG exchange, and as pathways for the spread of ARGs and human exposure. Next-generation sequence-based monitoring has recently enabled direct access and profiling of the total metagenomic DNA pool, where ARGs are identified or predicted based on the “best hits” of homology searches against existing databases. Unfortunately, this approach tends to produce high rates of false negatives. To address such limitations, we propose here a deep leaning approach, taking into account a dissimilarity matrix created using all known categories of ARGs. Two models, deepARG-SS and deepARG-LS, were constructed for short read sequences and full gene length sequences, respectively. Performance evaluation of the deep learning models over 30 classes of antibiotics demonstrates that the deepARG models can predict ARGs with both high precision (>0.97) and recall (>0.90) for most of the antibiotic resistance categories. The models show advantage over the traditional best hit approach by having consistently much lower false negative rates and thus higher overall recall (>0.9). As more data become available for under-represented antibiotic resistance categories, the deepARG models’ performance can be expected to be further enhanced due to the nature of the underlying neural networks. The deepARG models are available both in command line version and via a Web server at http://bench.cs.vt.edu/deeparg. Our newly developed ARG database, deepARG-DB, containing predicted ARGs with high confidence and high degree of manual curation, greatly expands the current ARG repository. DeepARG-DB can be downloaded freely to benefit community research and future development of antibiotic resistance-related resources.

**Abbreviations:**

ARG: antibiotic resistance gene

## INTRODUCTION

Antibiotic resistance is an urgent and growing global public health threat. It is estimated that the number of deaths due to antibiotic resistance will exceed ten million annually by 2050 and cost approximately 100 trillion USD worldwide (Jim, 2016; Brogan and Mossialos, 2016; O’Neill, 2014). Antibiotic resistance arises when microorganisms are able to survive an exposure to antibiotics that would normally kill them or stop their growth. This process allows for the emergence of ‘superbugs’ that currently are difficult to treat. A few examples include methicillin-resistant Staphylococcus aureus (MRSA), which is an extremely drug-resistant bacterium associated with several infections (Vuong et al., 2016), multidrug-resistant (MDR) Mycobacterium tuberculosis, which is resistant to rifampicin, fluoroquinolone, and isoniazid (Gandhi et al., 2010), and colistincarbapenem-resistant Escherichia coli, which have gained resistance to last-resort drugs through the acquisition of the MCR-1 and NDM-1 antibiotic resistance genes (ARGs) (Mediavilla et al., 2016; Hu et al., 2016).

The advent of high throughput DNA sequencing technology now provides a powerful tool to profile the full complement of DNA, including ARGs, derived from DNA extracts obtained from a wide range of environmental compartments, including livestock manure, compost, wastewater treatment plants, soil, water, and other affected environments (Pal et al., 2016; Forsberg et al., 2014; Berendonk et al., 2015; Pruden et al., 2013; Fahrenfeld et al., 2014; Mao et al., 2015), as well as within the human microbiome (Bengtsson-Palme et al., 2015; Pehrsson et al., 2016). Identification of ARGs from such samples is presently based on the computational principle of comparison of the metagenomic DNA sequences against available online databases. Such comparison is performed by aligning raw reads or predicted open reading frames (full gene length sequences) from assembled contigs to the database of choice, using programs such as BLAST (States and Agarwal, 1996), Bowtie (Langmead et al., 2009), or DIAMOND (Buchfink et al., 2015), and then predicting or assigning the types of ARGs present using a sequence similarity cutoff and sometimes an alignment length requirement (Yang et al., 2016; Zankari et al., 2012; Davis et al., 2016).

Existing bioinformatics tools focus on detecting known ARG sequences in genomic or metagenomic sequence libraries and thus are biased towards specific ARGs (McArthur and Tsang, 2016). For instance, ResFinder (Moran et al., 2016) and SEAR (Rowe et al., 2015) predict specifically plasmid-borne ARGs, and Mykrobe predictor (Bradley et al., 2015) is dedicated to 12 types of antimicrobials, while PATRIC (Davis et al., 2016) is limited to identifying carbapenem, methicillin, and beta lactam ARGs. Most of these tools use microbial resistance databases along with a “best hit” approach to predict whether a sequence is truly an ARG. Generally, predictions are restricted to high identity cutoffs, requiring a best hit with an identity greater than 80% by many programs such as ResFinder and ARGs-OAP (Pal et al., 2016; Yang et al., 2016; Zankari et al., 2012). In some studies the identity cutoff is even higher, as high as 90% for determining structure and diversity of ARGs through several resistomes (Pal et al., 2016) or analyzing the co-occurrence of environmental ARGs (Li et al., 2015).

Although the best hit approach has a low false positive rate, that is, few non-ARGs are predicted as ARGs (Forsberg et al., 2014), the false negative rate can be very high and a large number of actual ARGs end up being predicted as non-ARGs (Yang et al., 2016; McArthur and Tsang, 2016). Figure 1 shows the distribution of manually curated potential ARGs from the Universal Protein Resource (UNIPROT) database against the Comprehensive Antibiotic Resistance Database (CARD) and the Antibiotic Resistance Genes Database (ARDB). All of the gene comparisons indicate significant e-values < 1e-20 with the sequence identity ranging from 20% to 60% and bit scores > 50, which is considered statistically significant (Pearson, 2013). Thus, high identity cutoffs clearly will remove a considerable number of genes that are actual ARGs. For example, the entry <u>O07550 (Yhel),</u> a multidrug ARG conferring resistance to doxorubicin and mitoxantrone, has an identity of 32.47% with a significant e-value of 6e-77 to the best hit from the CARD database; the gene POCOZ1 (VraR), conferring resistance to vancomycin, has an identity of only 23.93% and an e-value 9e-13 to the best hit from the CARD database. Therefore, more moderate constraints on sequence similarity should be considered to avoid an unacceptable rate of false negatives. On the other hand, for short metagenomic sequences/reads (e.g., ~ 25aa or 100bp), a stricter identity constraint of ~80% is recommended (Pearson, 2013; Zankari et al., 2012) to avoid a high false positive rate. In principle, the best hit approach works well for detecting known and highly conserved types of ARGs but may fail to detect novel ARGs or those with low sequence identity to known ARGs (Yang et al., 2016; Xavier et al., 2016).

**Figure 1:**
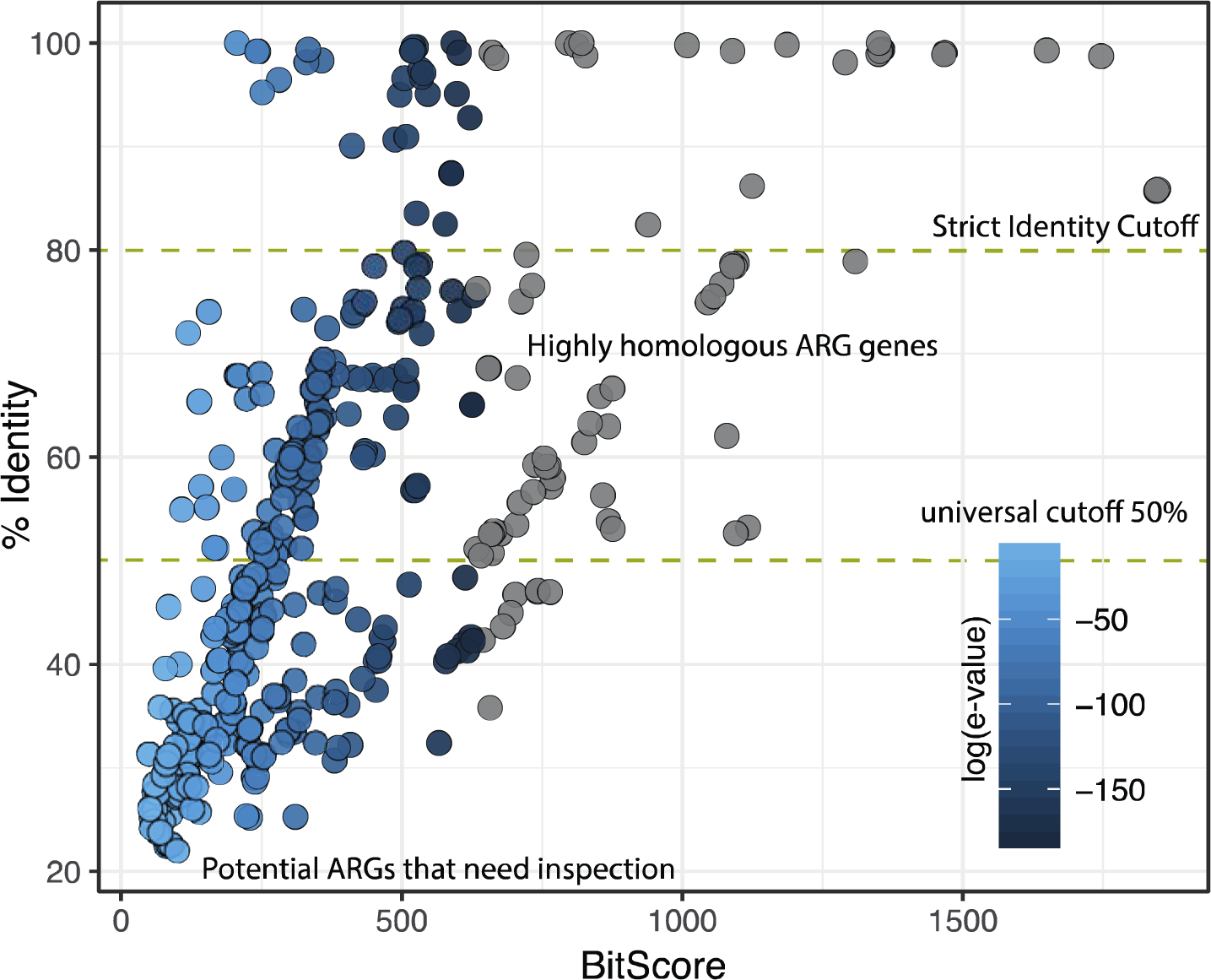
Bit score vs. identity distribution. This plot shows the relationship between the UNIPROT genes against the CARD and ARDB genes in terms of the percentage identity, bit score, and e-value. Colors depict the exponent of the e-value.

To address the limitation of current best hit methodologies, a deep learning approach was used to predict ARGs, taking into account the homology distribution of sequences in the ARG database, instead of only the best hit. Deep learning has proven to be the most powerful machine learning approach to date for many applications, including image processing (LeCun et al., 2015), biomedical signaling (Tabar and Halici, 2017), speech recognition (Hinton et al., 2012), and genomic-related problems, such as the identification of transcription factor binding sites in humans (Alipanahi et al., 2015; Pan and Shen, 2017). Particularly in the case of predicting DNA sequence affinities, the deep learning model surpasses all known binding site prediction approaches (Alipanahi et al., 2015). Here, we develop, train, and evaluate two deep learning models, deepARG-SS and deepARG-LS, to predict ARGs from short reads and full gene length sequences, respectively. The resulting database, deepARG-DB, is manually curated and contains predicted ARGs with a high degree of confidence, greatly expanding the current repertoire of ARGs currently accessible for metagenomic analysis of environmental datasets. DeepARG-DB can be queried either online or downloaded freely to benefit a wide community of users and to support future development of antibiotic resistance-related resources.

## METHODS

### DATABASE MERGING

The initial collection of ARGs was obtained from three major databases: CARD (Jia et al., 2017), ARDB (Liu and Pop, 2009) and UNIPROT (Apweiler et al., 2004). For UNIPROT, all genes that contained the Antibiotic Resistance keyword (KW-0046) were retrieved, together with their metadata descriptions when available. All identical or duplicate sequences were removed by clustering all the sequences (ARDB + CARD + UNIPROT) with CD-HIT (Huang et al., 2010), discarding all except one that have 100% identity and the same length. The remaining set of sequences comprises a total of 2,290 genes from ARDB (50% of the original ARDB genes), 2,161 from CARD (49% of the original CARD genes), and 28,108 from UNIPROT (70% of the original UNIPROT genes).

### ARG ANNOTATION IN ARDB AND CARD

The ARDB and CARD databases provide well-documented classification of ARGs, including the antibiotic class/type to which a gene confers resistance (e.g., macrolides, beta lactamases, or aminoglycosides) and the antibiotic group/subtype to which the gene belongs (e.g., tetA, sul1, macB, OXA, MIR, or DHA). Manual inspection revealed that some genes have been assigned to specific sets of antibiotics instead of antibiotic resistance classes or categories. For instance, carbapenem, carbenicillin, cefoxitin, ceftazidime, ceftriaxone, and cephalosporin are actually a subset of the beta lactamases class. Thus, a total of 102 antibiotics that were found in the ARDB and CARD databases were further consolidated into 30 antibiotic categories (see supplementary Table S1).

## UNIPROT GENE ANNOTATION

Compared to the ARGs in CARD and ARDB, the UNIPROT genes with antibiotic resistance keywords are less well curated. Therefore, additional procedures were applied to further annotate the UNIPROT genes. Specifically, based on the CD-hit (Huang et al., 2010) clustering results, clusters that contained only UNIPROT genes were classified into two categories: 1) those without any annotation were tagged as “unknown” and 2) those with descriptions were text mined to identify possible association with antibiotic resistance.

UNIPROT’s sequence description contains a variety of features including a description of possible functions of the protein, the gene name containing the HUGO nomenclature (Eyre et al., 2006) for each sequence, and the evidence indicating whether a sequence has been manually inspected or not. A text mining approach was used to mine the genes’ descriptive features to identify their antibiotic resistance associations with the 102 antibiotic names and 30 antibiotic classes. The Levenshtein distance (Yujian and Bo, 2007) was used to measure the similarities between gene description and antibiotic categories. This text mining approach was used because the names of the antibiotic resistance categories are not standardized among the databases and flexibility is needed to identify as many antibiotic associations as possible. For instance, genes linked to beta lactamases were sometimes tagged as beta-lactam, beta-lactamases, or beta-lactamase. Thus, text mining using all the alternative words allows comprehensive identification of antibiotic associations for each gene. Using this strategy, genes from UNIPROT were tagged either to their antibiotic resistance associations based on their description, or to “unknown” if no link to any antibiotic was found. Then manual inspection was performed to remove misleading associations that passed the similarity criteria. The final set of genes and their tagged antibiotic resistance classes or categories are shown in Figure 2. Altogether, 16,360 UNIPROT genes remained after this refinement procedure.

**Figure 2:**
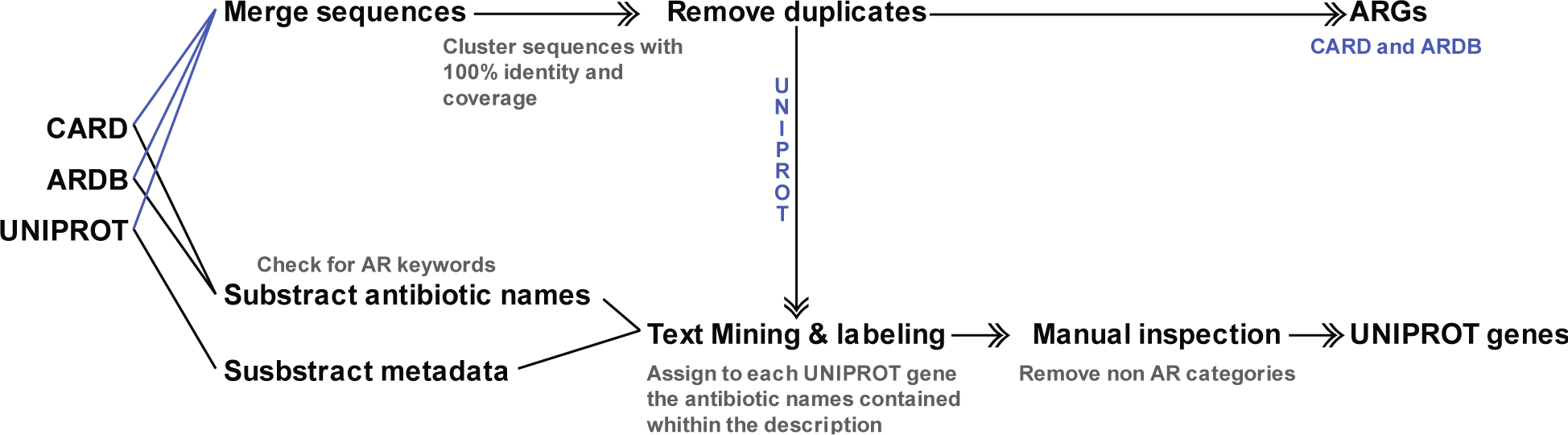
Preprocessing and UNIPROT ARGs annotation. Antibiotic resistance genes from CARD, ARDB, and UNIPROT are merged and clustered to remove duplicates. Then sequences from UNIPROT are annotated using the matches between the metadata and the antibiotic names from ARDB and CARD.

The text mining procedure enabled the UNIPROT genes to become linked to one or more classes of antibiotics. However, the text mining procedure is purely based on gene metadata. Therefore, there was no evidence at the sequence level that the UNIPROT genes were truly associated with antibiotic resistance. For that reason, the UNIPROT gene’s annotation was further validated by their sequence homology to the CARD and ARDB databases. DIAMOND, a program that has similar performance to BLAST (Healy, 2007), but is much faster (Buchfink et al., 2015), was used for this purpose. For simplicity, UNI-gene is used here to denote a UNIPROT-derived gene, and CARD/ARDB-ARG is used to denote a gene derived from either CARD or ARDB (Figure 3). According to the sequence homology, each UNI-gene was classified into the following categories based on their potential to confer antibiotic resistance (defined as annotation factor):

1. High quality ARGs (High): A UNI-gene is tagged with a “High” annotation factor if it has ≥ 90% identity to a CARD/ARDB-ARG over its entire length. This similarity cutoff has been used in other studies to identify relevant ARGs (Pal et al., 2015; Li et al., 2017) and is stricter than that used in the construction of the ARDB database (Liu and Pop, 2009).
2. Homologous ARGs (Mid): A UNI-gene is tagged with a “Mid” annotation factor if it has ≥ 50% identity and an e-value lower than 1e-10 to a CARD/ARDB-ARG and also consistent annotation to the CARD/ARDB-ARG.
3. Potential ARGs (Manual Inspection): A UNI-gene is tagged with “Manual inspection” if it has < 50% identity and an e-value lower than 1e-10 to CARD/ARDB-ARGs and also consistent annotation to CARD/ARDB-ARGs. This gene is considered a potential ARG but with insufficient evidence and therefore warrants further analysis for the veracity of its antibiotic resistance.
4. Discarded ARGs (Low): A UNI-gene is discarded if its annotation differs from the best hit CARD/ARDB-ARG and the e-value is greater than 1e-10. Note the gene can potentially still be an ARG, but due to a lack of sufficient evidence, it is removed from current consideration to ensure ARG annotation quality.

**Figure 3:**
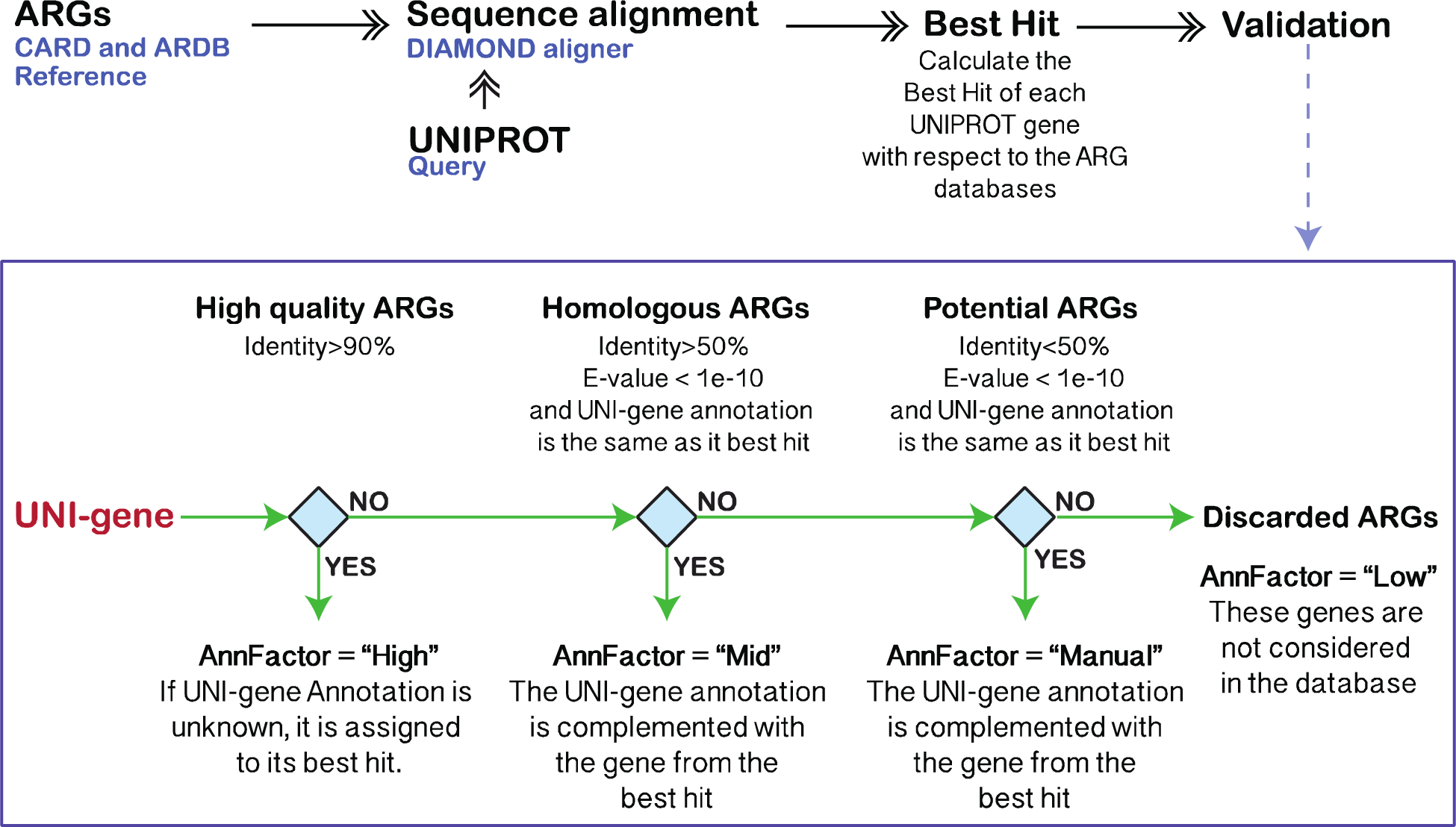
Validation of UNIPROT annotations. UNIPROT genes are aligned against the CARD and ARDB databases. The alignment with the highest bit score is selected for each UNI-gene (best hit) and a set of filters are applied to determine the UNI-gene annotation factor (AnnFactor).

Altogether 16,222 genes were tagged in the categories of “High” and “Mid” annotation factors. After removing sequences annotated as conferring resistance by single nucleotide polymorphisms (SNPs), a total of 14,933 genes were remaining for downstream construction of the deep learning models.

## DEEP LEARNING

Supervised machine learning models are usually divided into characterization, training, and prediction units. The characterization unit is responsible for the representation of DNA sequences as numerical values called features. It requires a set of DNA descriptors that are based on global or local sequence properties. Here the concept of dissimilarity based classification (Sorensen et al., 2010) is used, where sequences are represented and featured by their homology distances to known ARGs. The CARD and ARDB genes are chosen for the known ARGs whereas the UNIPROT (High+Mid) genes are used for training and validation of the models (see Figure 4). The bit score was used as the similarity indicator, because it takes into account the extent of homology between sequences and, unlike the e-value, is independent of the database size (Pearson, 2013). The process for computing the dissimilarity representation was carried out as follows. The UNIPROT genes were aligned to the CARD and ARDB databases (Jia et al., 2017; Liu and Pop, 2009) using DIAMOND (Buchfink et al., 2015) with very permissive constraints: 10,000 maximum number of hits representing the total number of reported hits to which a UNIPROT gene is aligned, a 20% minimum identity (--id 20), and an e-value smaller than 1e-10. The bit score was then scaled to the [0, 1] interval, where 0 represents a statistically significant alignment and 1 a lack of sequence alignment. Thus, a feature matrix was built where the rows correspond to the homology similarity of the UNIPROT genes to the features (ARDB/CARD genes).

**Figure 4:**
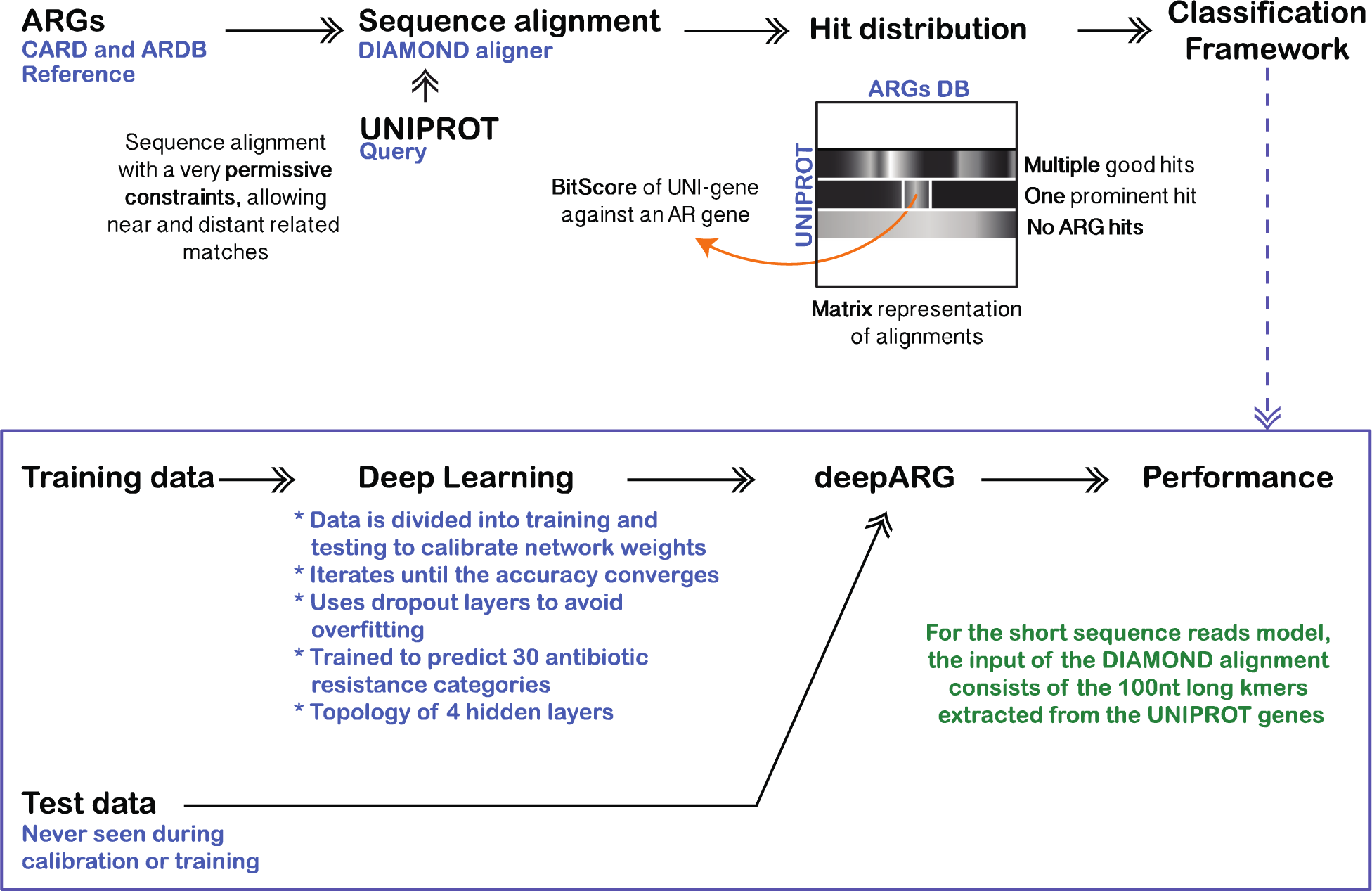
Classification framework. UNIPROT genes are used for validation and training whereas the CARD and ARDB databases are used as features. The distance between genes from UNIPROT to ARGs databases is computed using the sequence alignment bit score. Alignments are done using DIAMOND with permissive cutoffs allowing a high number of hits for each UNIPROT gene. This distribution is used to train and validate the deep learning models.

A deep learning model, deepARG, was then created to annotate metagenomic sequences to antibiotic resistance categories. One of the main advantages of deep learning over other machine learning techniques is its ability to discriminate relevant features without the need for human intervention (Min et al., 2016; Coates et al., 2011; Sun et al., 2014). It has been highlighted for its good performance to resolve multiclass classification problems (Buggenthin et al., 2017; Alipanahi et al., 2015; Qin and Feng, 2017; Dong et al., 2016; Bhatkoti and Paul, 2016). Here, a deep learning multiclass model was trained by taking into account the homology distance distribution of a sequence to all known ARGs. This distribution represents a high level of sequence abstraction propagated through a fully connected network. The deepARG model consists of four dense hidden layers of 2000, 1000, 500, and 100 units that propagate the bit score distribution to dense and abstract features. The input layer consists of 4,333 units that correspond to the ARGs from ARDB and CARD. These features are used during training and evaluation. To avoid overfitting, random hidden units were removed from the model at different rates using the dropout technique (Baldi and Sadowski, 2014). Lastly, the output layer of the deep neural network consists of 30 units that correspond to the antibiotic resistance categories (see Supplementary Table 1). The output layer uses a softMax (Dunne and Campbell, 1997; Sundermeyer et al., 2012) activation function that computes the probability of the input sequence against each ARG category. The probability is used to define the ARG category to which the input sequence belongs. The deepARG architecture is implemented using the Python Lasagne (Van Merriënboer et al., 2015) module, a high level wrapper for the widely used Theano (Bergstra et al., 2011) deep learning library. Because deep learning demands intensive computational resources, the training was carried out using the GPU routines from Theano. However, heavy computation was required only once to obtain the deep learning model and the prediction routines do not require such computational resources. The deepARG architecture has been successfully tested on a laptop with 8Gb RAM and i7 processor.

Two strategies have been used to identify ARGs in metagenomic data; one predicts ARGs directly using short reads, while the other uses predicted open reading frames (i.e., full gene length sequences) from assembled contigs to predict ARGs. To allow for both annotation strategies, two deep learning models, deepARG-SS and deepARG-LS, were developed to process short reads and full gene length sequences, respectively. The deepARG-SS model was designed specially to classify short reads generated by NGS technologies such as Illumina. DeepARG-LS was trained using complete ARG sequences and can be used to annotate novel ARG genes, for instance, in open reading frames detected in assembled contigs from the MetaHit consortium (Qin et al., 2010). Note that each model was trained and validated separately to ensure high performance.

To evaluate the performance of the deepARG models (deepARG-SS and deepARG-LS), two strategies were used. First, 70% of the ARGs from UNIPROT were used as training and the remaining 30% for validation. Second, MEGARes (Lakin et al., 2017), a recently developed database containing manually curated ARGs from CARD (Jia et al., 2017), ARG-ANNOT (Gupta et al., 2014), and RESFINDER (Zankari et al., 2012), was used as an independent dataset to further check the performance of the deepARG-LS model. In addition, the performance of the deepARG models was compared with the best hit approach with a minimum identity cutoff of 80%. The prediction quality was evaluated by precision, recall, and F1-score metrics defined as,

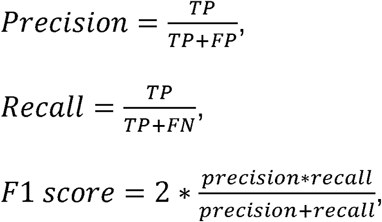

where TP represents true positives (i.e., an ARG is predicted correctly as an ARG), FP false positives (a nonARG is predicted as an ARG), and FN false negatives (an ARG is predicted as a nonARG).

Note because the first step of the deepARG pipeline consists of the sequence alignment using DIAMOND, nonARGs (short reads or full length genes) are filtered out and not considered for further prediction. Therefore, the alignment stage only passes ARG-like sequences that have e-value<1e-10 and identity>20% to deepARG for prediction. Thus, the performance reflects the capability of the deepARG models in differentiating the 30 antibiotic resistance categories.

## RESULTS AND DISCUSSION

After the databases were merged and duplicates and SNPs were removed, a total of 2,203, 2,130, and 28,108 genes were collected from the ARDB (50% of full ARDB), CARD (49% of all CARD genes), and UNIPROT (70% of total ARG-like sequences from UNIPROT) databases, respectively. For UNIPROT genes, a total of 16,360 genes were annotated using the available gene description. Following validation through sequence homology, 10,602 ARG-like sequences remained. The resulting database, deepARG-DB, comprises 30 antibiotic classes/types, 2,149 subtypes, and 14,933 reference sequences. Over 32% of the genes belong to the Beta lactamase class (5136), followed by 26% the bacitracin class (4205), 7.7% macrolide-lincosamide-streptogramin (MLS) (1,109), 5.6% aminoglycoside (915), 5.4% polymixin (879) and 5.4% multidrug (877), accounting for the major antibiotic resistance classes in deepARG-DB (see Figure 6A). The categories where the UNIPROT database made the greatest contribution correspond to beta-lactam, bacitracin, MLS, and polymyxin. However, not all ARG categories were found in the UNIPROT database, such as elfamycin, fusidic acid, puromycin, among others (see Figure 6B for details). One of the limitations of deepARG-DB is its dependency on the quality of the CARD and ARDB databases. Thus, to avoid the propagation of errors from the CARD and ARDB, gene categories and groups were manually inspected and corrected. In particular, those annotations that differ from the ARDB and CARD databases. Because UNIPROT and CARD are constantly updated, the deepARG-DB will get updated and versioned as well as the trained deep learning models.

**Figure 5:**
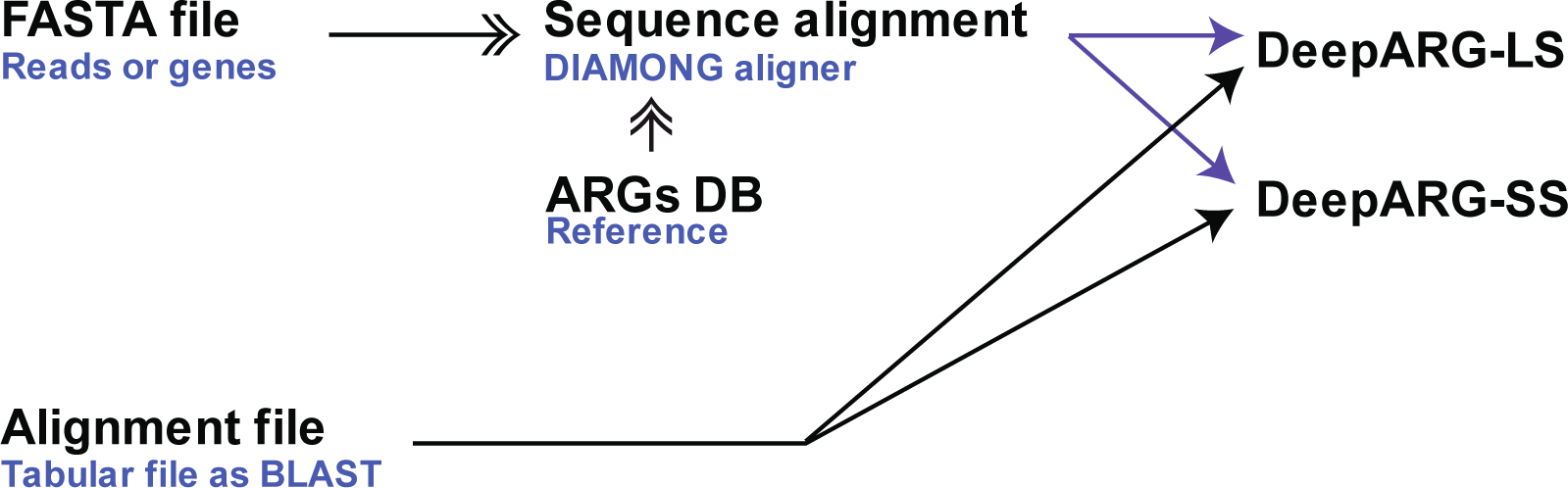
deepARG pipeline. Input can be short reads or full gene-like sequences. Users can upload the sequences in FASTA format or the sequence alignments from any program that reports the bit score as BLAST (e.g., DIAMOND). Then, sequences are annotated using one of the deep learning models.

**Figure 6:**
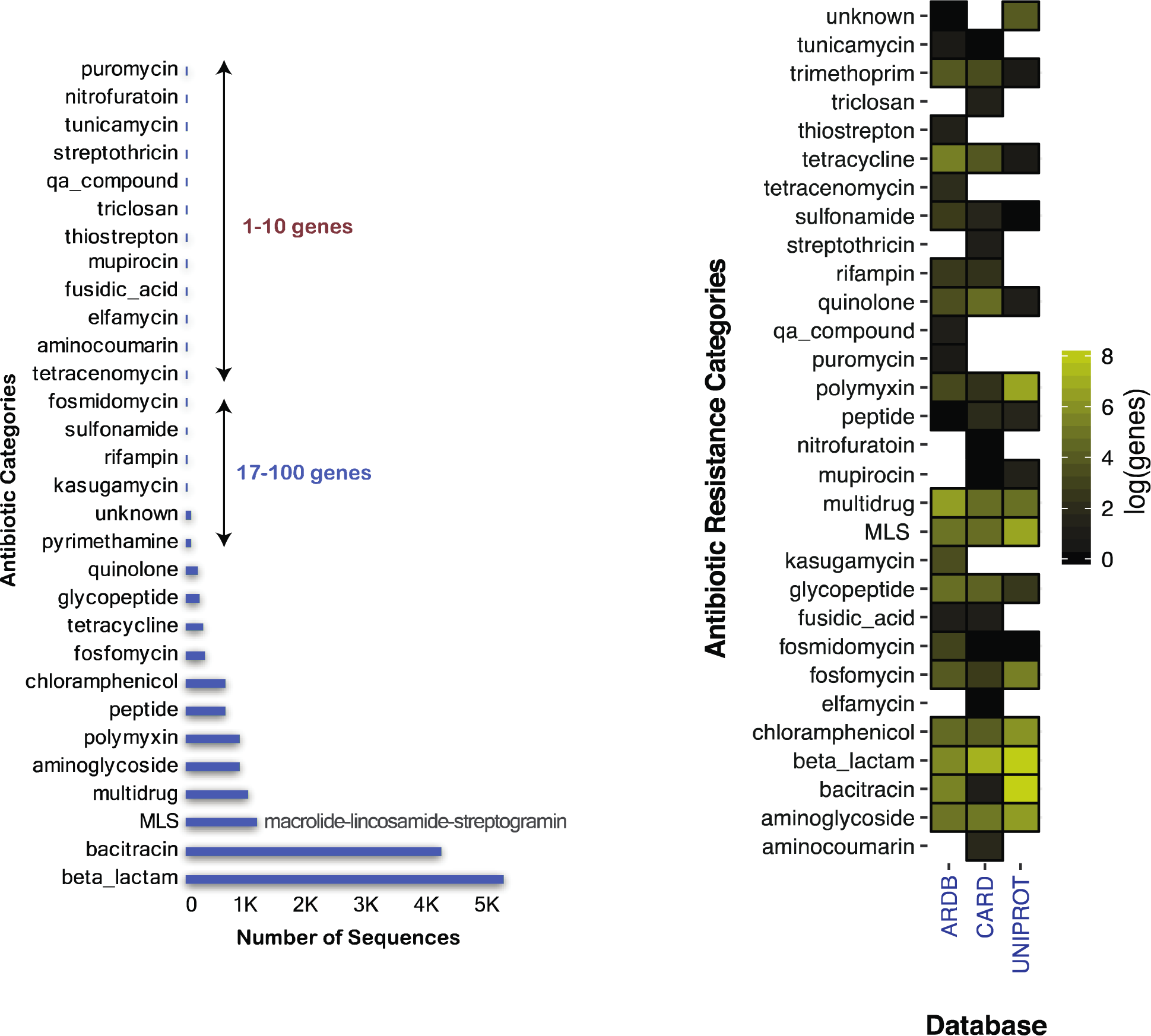
A) Distribution of the number of sequences in the 30 antibiotic categories in deepARG-DB. B) The relative contribution of ARG categories in the ARDB, CARD, and UNIPROT databases.

### PREDICTION OF SHORT SEQUENCE READS

To simulate a typical metagenomic library, UNIPROT genes were split into 100 nucleotide long sequences, with a total of 321,008 reads generated. The deepARG-SS model was subsequently trained and tested in a manner in which 70% of the reads were randomly selected for training, while the remaining 30% were reserved for validation. An overall precision of 0.97 and a recall of 0.91 were achieved among the 30 antibiotic categories tested (see Figure 7A). In comparison, the best hit approach achieved an overall 0.96 precision and 0.51 recall. Achieving high precision for the best hit approach is not surprising, as the method relies on high identity constraints and has been reported to predict a low number of false positives, but a high number of false negatives (Yang et al., 2016). We observed that both methods yielded high precision for most of the categories (see Figure 7B). However, both methods performed poorly for the triclosan category, which was populated only by four genes.

**Figure 7:**
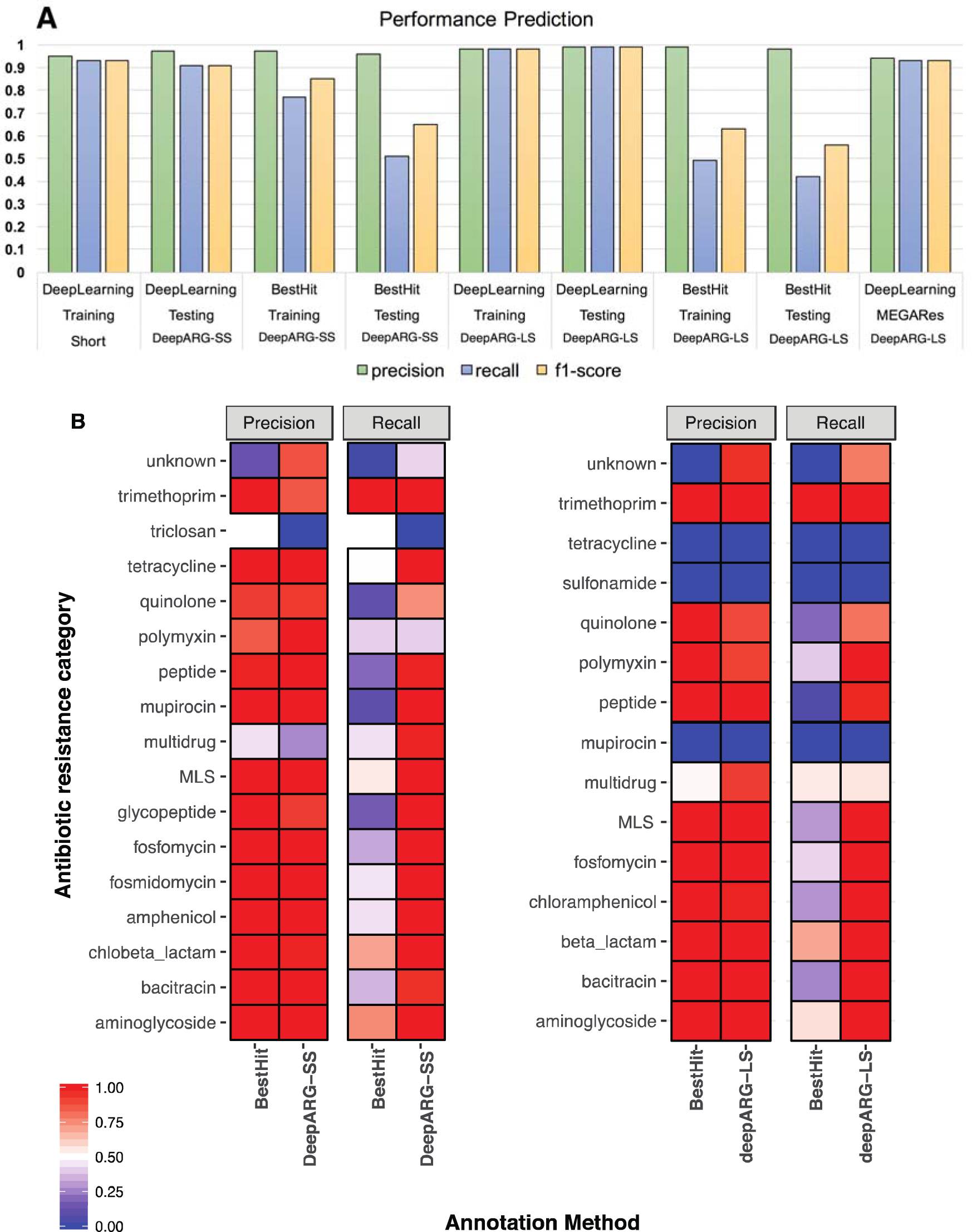
A) Performance comparison of the deepARG models with the best hit approach using precision, recall, and F1-score as metrics for the training and testing datasets. The MEGARes bars corresponds to the performance of deepARG-LS using the genes from the MEGARes database. B) Precision and recall of deepARG models against the best hit approach for each individual category in the testing dataset. *UNIPROT genes are used for testing and not all the ARG categories have genes from the UNIPROT database.

The deepARG-SS model performs particularly well for antibiotic resistance categories containing a large number of ARGs, such as beta lactamases, bacitracin, and MLS, but not as well for categories with a small number of ARGs, such as triclosan and aminocoumarin. This result is expected due to the nature of neural network models. As more data becomes available to train the models, the better their ultimate performance. In contrast, the best hit approach yields perfect prediction for some ARG categories containing a limited number of ARGs, but not for categories with a large number of ARGs (see Figure 7B and supplementary Table S2 for details).

For the multidrug antibiotic resistance category, the deepARG-SS model had an almost perfect recall (0.99), implying that only a small number of multidrug reads were classified to other categories. However, the deepARG-SS model also had the highest false positive rate compared to other classes (precision 0.27), implying that many non-multidrug reads were annotated as multidrug sequences. On the other hand, the best hit approach showed a higher precision (0.44), but a much lower recall (0.44). The multidrug category contains genes that confer resistance to multiple antibiotic classes such as macrolides, beta-lactamases, glycopeptides, quinolones, as well as other antimicrobials such as metals (Chen et al., 2017; Linhares et al., 2015). These genes often share similar sequences, which makes it challenging for computational methods to determine the true source of a short read. Therefore, when reads yield a best prediction probability less than 0.9, deepARG reports the top two ARG categories for further examination. Since the low precision seen in both methods suggests that other non-multidrug categories may contain genes that are close to the multidrug category, it is clear that there is still much room for improvement in the existing annotation of ARGs.

Contrary to the multidrug category, the “unknown” antibiotic resistance category has a high precision of 0.87, but a low recall of 0.42, indicating a high false negative rate. Thus, reads from the unknown antibiotic resistance class are mistakenly assigned/predicted to other antibiotic resistance categories. This highlights the need to check whether the unknown category actually contains genes from other ARG classes such as beta-lactam, macrolides, triclosan, among others. Comparatively, the best hit approach has the worst performance for the “unknown” antibiotic class (see Figure 7B and supplementary Table S2). In general, the deepARG-SS model shows a significant improvement over the false negative rate compared to the best hit approach for almost all ARG categories.

### PREDICTION OF LONG GENE-LIKE SEQUENCES

The deepARG-LS model was trained and tested using full gene-length sequences. The UNIPROT validated genes were split into a training set (70% of the data) and a validation set (30% of the data). The CARD and ARDB databases were used as features. The deepARG-LS model shows similar results, yielding an overall 0.99 precision and 0.99 recall for predicting different classes of ARGs. Better performance in deepARG-LS than deepARG-SS is expected because longer sequences contain more information than short reads (Figure 7). Particularly deepARG-LS achieved a high precision (0.97 ± 0.03) and a perfect recall (0.99 ± 0.01) for the antibiotic classes that are highly represented, such as bacitracin, beta lactamase, chloramphenicol, and aminoglycoside (See Figure 7B and supplementary Table S3 for details). Comparatively, the best hit approach achieved a perfect precision (1.00 ± 0.00) but a much lower recall (0.48 ± 0.2) for these classes. Similar to deepARG-SS, deepARG-LS did not perform well for categories with few genes, such as sulfonamide and mupirocin (See supplementary Table S3 for details).

### PERFORMANCE PREDICTION OF KNOWN AND VALIDATED ARGS

To further evaluate and validate performance, the deepARG-LS model was applied to all of the ARG sequences in the MEGARes database, which excludes resistance mechanisms due to SNPs (Lakin et al., 2017). Comparison of the deepARG-LS prediction with the database annotation yielded an overall precision and recall of 0.94 and 0.93, respectively (Figure 7A and supplementary Table S4). The deepARG-LS model achieved a perfect precision of 0.99 ± 0.05 and recall of 0.96 ± 0.03 for categories with a large number of genes, such as beta lactamases, elfamycin, fosfomycin, glycopeptides, MLS, and sulfonamide. However, the model performed poorly for the classes that have a small number of genes (see Supplementary Table S4). For instance, MEGARes has a Tunicamycin gene that was labeled by the deepARG-LS model as quinolone with a probability of 0.6. The low probability 0.6 suggests that the gene has more than one annotation. When the complete annotation for this gene was inspected, it was found that the deepARG-LS model predicted the correct label (Tunicamycin) with a 0.3 probability, indicating that for this particular class more gene sequences are needed for the training of the model. The deepARG-DB database has only three Tunicamycin genes, which may explain why this gene was not properly classified. However, it is worth noting that the thiostrepton category was predicted correctly despite its lower number of training genes. The multidrug category is one of the most difficult categories to predict. It contains about 200 genes and the deepARG-LS model yielded a 0.7 precision with a 0.6 recall. This result suggests the need to reexamine the genes tagged as multidrug as well as the genes from other categories that were assigned to the multidrug class. Overall, it also highlights the need to review different antibiotic resistance databases to find a consensus about what is considered to be a multidrug resistance gene.

### STANDALONE DeepARG PROGRAM AND THE WEB SERVER

The source code for the deepARG models can be downloaded from the git repository (https://gaarangoa@bitbucket.org/gaarangoa/deeparg-ss.git). It consists of a command line program where the input can be either a FASTA file or a BLAST tabular file. If the input is a FASTA sequence file, deepARG will perform the sequence search first and then annotate ARGs. If the input is already a BLAST tabular file, deepARG will annotate ARGs directly (see Figure 5 for details). An online version of deepARG is also available where a user can upload a FASTA file of the sequences for ARG annotation (http://bench.cs.vt.edu/deeparg). Once the data is processed, the user receives an email with results of annotated ARGs and absolute abundance of the ARGs. Using the command line version, the user also has access to more elaborated results such as the probabilities of each read/gene belonging to the specific antibiotic resistance categories. In addition to prediction of antibiotic classes and the associated probabilities, the deepARG model reports the entries with multiple classifications. In detail, if a read or complete gene sequence is classified to an antibiotic category with a probability below 0.9, the top two classifications will be provided. This would help researchers identify reads/sequences with less confident predictions, and it is recommended that the detailed output be examined together with domain knowledge to determine the more likely ARG category. The DeepARG-DB is freely available under the DeepARG Web site (http://bench.cs.vt.edu/deeparg) as a protein FASTA file. Each entry in the database has a complete description that includes the gene identifier, the database where the gene is coming from, the antibiotic class and the antibiotic group.

## CONCLUSIONS

Here a new computational resource for the identification and annotation of ARGs derived from metagenomic data is developed, trained, and evaluated. The deep learning approach proved to be more accurate than the widely used best hit approach and is not restricted to strict cutoffs, thus greatly reducing false negatives and offering a powerful approach for metagenomic profiling of ARGs in environmental compartments. Further, the deepARG database developed here greatly expands the available ARGs individually available in the currently most widely used CARD, ARDB, and UNIPROT databases, including their existing sequence content and extensive metadata. DeepARG provides a publicly available database structured into a simple class and group hierarchy for each ARG. While deepARG is not intended to replace CARD or ARDB, in conjunction with deep learning, it aims to improve the ARG annotation by drastically reducing the false negative rate, while maintaining a similarly high true positive rate associated with the traditional best hit approach.

## ACKNOWLEDGEMENT

Funding was provided in part for this effort by USDA NIFA Award # Effective Mitigation Strategies for Antimicrobial Resistance program, the NSF Partnership in International Research and Education (PIRE) award #1545756, the Virginia Tech Institute for Critical Technology and Applied Science Center for the Science and Engineering of the Exposome (SEE) and the Virginia Tech Sustainable Nanotechnology Interdisciplinary Graduate Education Program (IGEP).

